# Foreground-background segmentation revealed during natural image viewing

**DOI:** 10.1101/109496

**Authors:** Paolo Papale, Andrea Leo, Luca Cecchetti, Giacomo Handjaras, Kendrick Kay, Pietro Pietrini, Emiliano Ricciardi

**Affiliations:** Molecular Mind Lab, IMT School for Advanced Studies Lucca, Lucca, 55100 Italy; Center for Magnetic Resonance Research, Department of Radiology, University of Minnesota, Twin Cities, Minneapolis, MN, 55455, USA

## Abstract

One of the major challenges in visual neuroscience is represented by foreground-background segmentation. Data from nonhuman primates show that segmentation leads to two distinct, but associated processes: the enhancement of neural activity during figure processing (i.e., foreground enhancement) and the suppression of background-related activity (i.e., background suppression). To study foreground-background segmentation in ecological conditions, we introduce a novel method based on parametric modulation of low-level image properties followed by application of simple computational image-processing models. By correlating the outcome of this procedure with human fMRI activity measured during passive viewing of 334 natural images, we reconstruct easily interpretable “neural images” from seven visual areas: V1, V2, V3, V3A, V3B, V4 and LOC. Results show evidence of foreground enhancement for all tested regions, while background suppression specifically occurs in V4 and LOC. “Neural images” reconstructed from V4 and LOC revealed a preserved spatial resolution of foreground textures, indicating a richer representation of the salient part of natural images, rather than a simplistic model of object shape. Our results indicate that scene segmentation is an automatic process that occurs during natural viewing, even when individuals are not required to perform any particular task.

## Introduction

In the scientific journey toward a satisfying understanding of the human visual system, scene segmentation represents a central problem “for which no theoretical solution exists” (Wu MC et al. 2006). Segmentation into foreground and background is crucial to make sense of the surrounding visual environment, and its pivotal role as an initial step of visual content identification has long been theorized (Biederman I 1987). Indeed, according to Fowlkes and colleagues (2007), humans can produce consistent segmentations of natural images. However, even though more recent approaches based on deep convolutional networks produced promising results (He K et al. 2017), both the computational and neurophysiological processes that underlies scene segmentation are still a matter of debate.

To date, numerous studies found evidence of texture segmentation and figure-ground organization in the early visual cortex of nonhuman primates (Lamme VA 1995; Lee TS et al. 1998; Poort J et al. 2012; Self MW et al. 2013) and humans (Kastner S et al. 2000; Scholte HS et al. 2008; Kok P and FP de Lange 2014). In particular, a recent study on monkeys attending artificial stimuli revealed an early enhancement of V1 and V4 neurons when their receptive fields covered the foreground, and a later response suppression when their receptive fields were located in the stimulus background (Poort J et al. 2016). This demonstrates that foreground enhancement and background suppression are distinct but associated processes involved in segmentation.

In addition, from an experimental viewpoint, the role of visual segmentation has been demonstrated only by means of non-ecological stimuli (e.g., binary figures, random dots, oriented line segments and textures). Although two recent studies investigated border-ownership in monkeys with both artificial and natural stimuli (Hesse JK and DY Tsao 2016; Williford JR and R von der Heydt 2016), a proof of the occurrence of foreground-background segmentation in the human brain during visual processing of naturalistic stimuli (e.g., natural images and movies) is still lacking.

In light of this, we specifically investigated foreground enhancement and background suppression, as specific processes involved in scene segmentation, during passive viewing of natural images. We used fMRI data, previously published by Kay and colleagues (Kay KN et al. 2008), to study brain activity from seven visual regions of interest (ROIs): V1, V2, V3, V3A, V3B, V4 and lateral occipital complex (LOC) during the passive perception of 334 natural images, whose “ground-truth” segmented counterparts have been included in the Berkeley Segmentation Dataset (BSD) (Arbelaez P et al. 2011).

Notwithstanding, as a reliable description of computational and neurophysiological processes involved in scene segmentation has not been achieved yet, we developed a novel pre-filtering modeling approach to study brain responses to complex, natural images without relying on explicit models of scene segmentation, and adopting a validated and biologically plausible description of activity in visual cortices. Our method is similar to other approaches where explicit computations are performed on representational features rather than on the original stimuli (Naselaris T et al. 2011). For instance, these methods have been recently used to investigate semantic representation (e.g. Huth AG et al. 2012; Handjaras G et al. 2016) or boundary and surface-related features (Lescroart M et al. 2016). However, as opposed to the standard modeling framework – according to which alternative models are computed from the stimuli to predict brain responses – here, low-level features of the stimuli are parametrically modulated and simple descriptors of each filtered image (e.g., edges position, size and orientation) are aggregated in a fixed model (Figure 1). The correspondence between the fixed model and fMRI patterns evoked by the intact images, was then assessed using representational similarity analysis (RSA) (Kriegeskorte N et al. 2008). Notably, this approach can also be exploited to obtain highly informative “neural images” representing the putative computations of different brain regions and may be generalized to investigate different phenomena in visual neuroscience.

**Figure 1.**
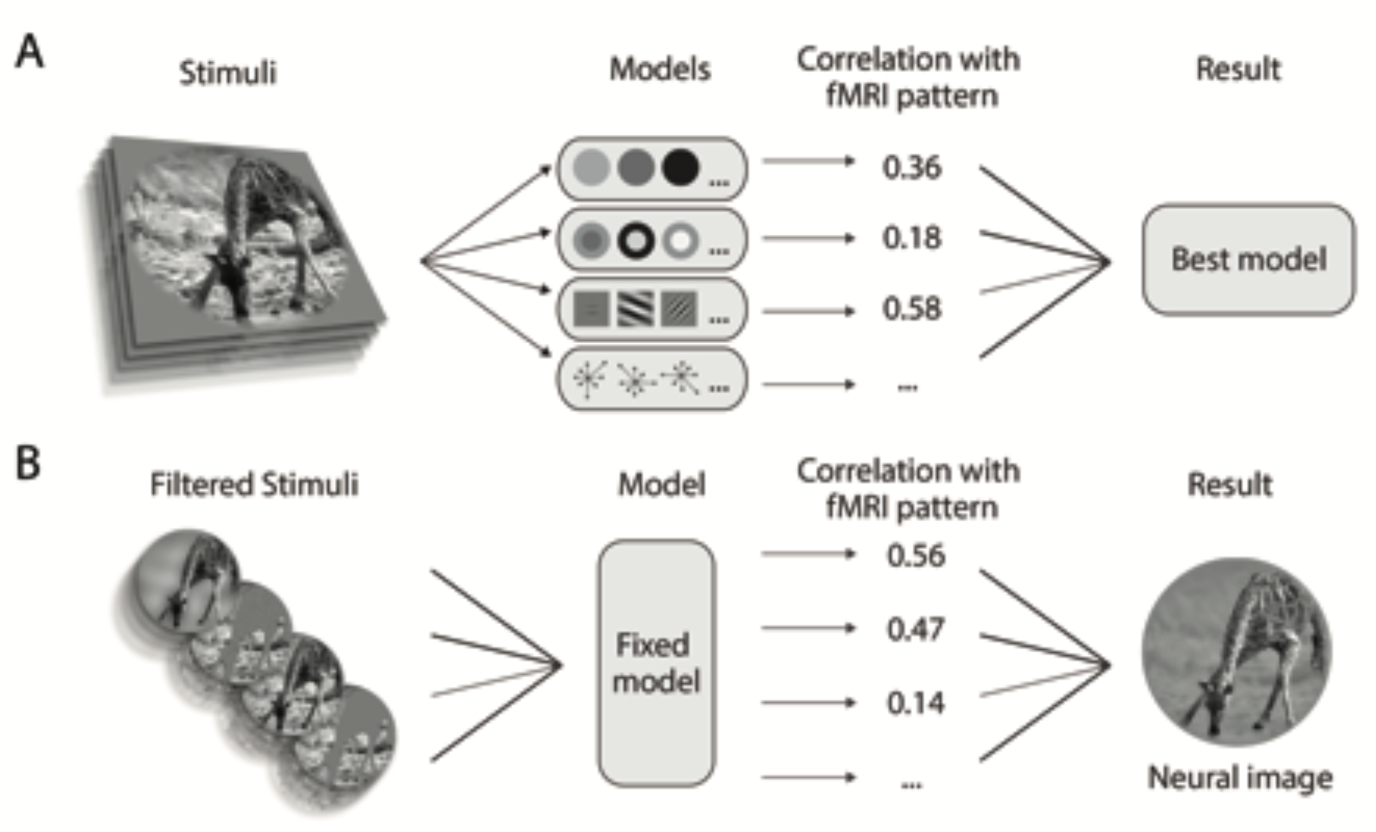
Comparing the Standard Modeling Approach and the Pre-Filtering Modeling Approach. A) In the standard modeling pipeline, different models are compared. After extracting features from the stimuli, competing feature vectors can be used in order to predict brain activity in an encoding procedure, or stimuli dissimilarities can be used in a representational similarity analysis. Finally, the model that better predicts brain responses is discussed. B) In our pre-filtering modeling approach, different filtered versions of the original stimuli are compared. Various biologically plausible filtering procedures are applied to the stimuli prior to compute a unique feature space according to a given fixed and easily interpretable model. In our approach a single model is employed and the best step of each filtering procedure is used to build a *post-hoc* “neural image”, to visually interpret the results. While the standard modeling approach is theoretically more advantageous, as its output is a fully computable model of brain activity, it can not be applied when reliable explicit models of the perceptual process do not exist yet, as in the case of scene segmentation. Alternative attempts to reconstruct visual stimuli from brain activity have been previously reported using decoding techniques (e.g. Stanley GB et al. 1999; Thirion B et al. 2006; Miyawaki Y et al. 2008; Nishimoto S et al. 2011).

To summarize, by examining the correlation between biologically plausible features extracted from manipulated stimuli and brain activity of different visual cortices during passive perception of natural intact images, we believe that new and relevant information on scene segmentation processing can be inferred.

## Materials and Methods

To assess differences between cortical processes involved in foreground-background segmentation, we employed a low-level description of images, defined by averaging the representational dissimilarity matrices (RDMs) of four well-known computational models (Figure 2D). These models are based on simple features – such as edge position, size and orientation – whose physiological counterparts are well known (Marr D 1982). The averaged model was kept constant while the images were parametrically filtered and iteratively correlated with brain activity through RSA. For each ROI, this pre-filtering modeling approach led to a pictorial and easily interpretable representation of the optimal features (contrast and spatial frequencies) of foreground and background of natural images (i.e., “neural images”). The analytical pipeline is schematized in Figure 2.

**Figure 2.**
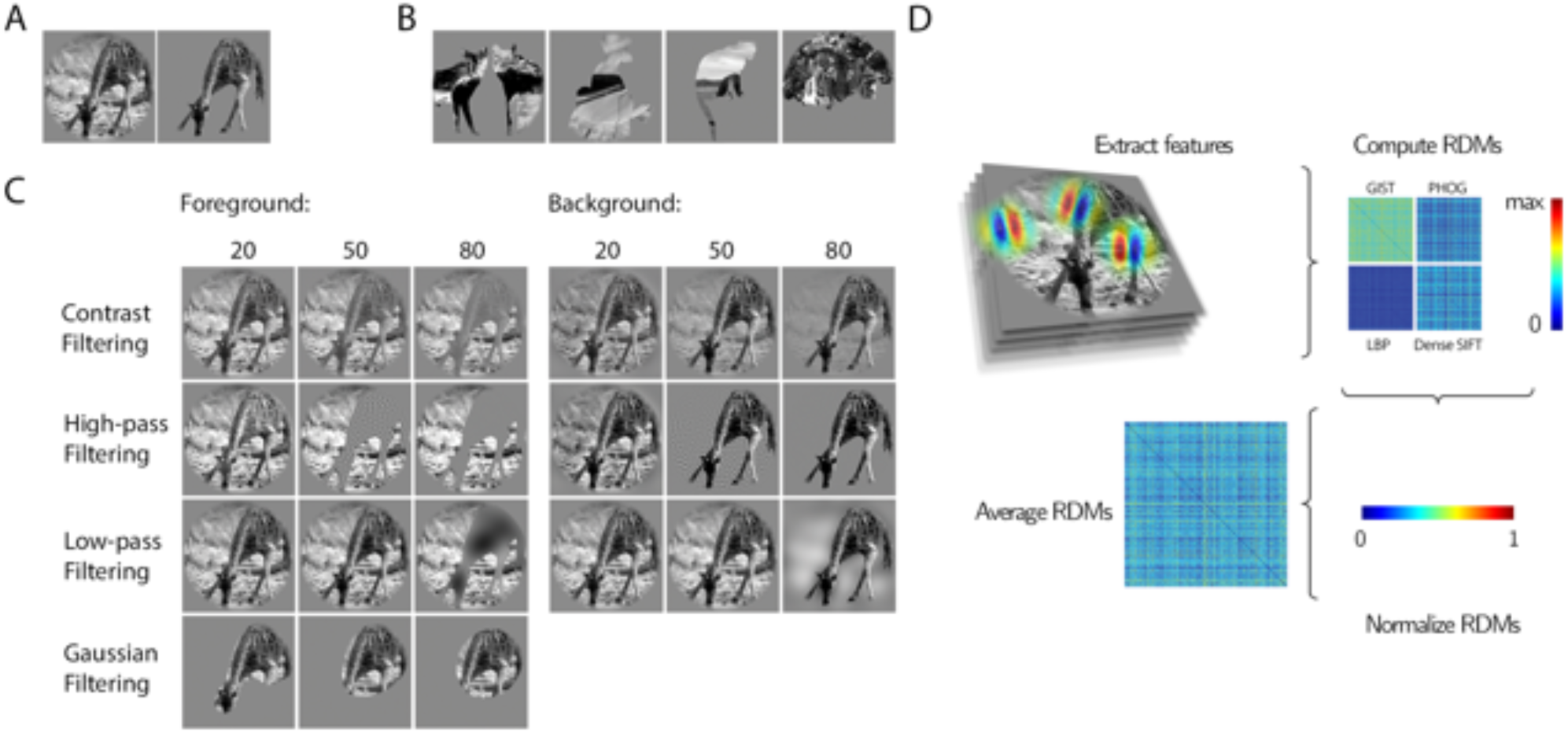
Analytical Pipeline. A) An example of intact image and its behaviorally segmented counterparts B) The set of segmented stimuli is tested against a null distribution of 1,000 permutations. Each permutation is built by randomly shuffling the 334 behavioral foreground masks C) Three steps (20, 50 and 80 out of 100) for the contrast, Gaussian or spatial frequencies filtering. D) In clockwise order: features for each model were extracted from the stimuli; the dissimilarity (1- Pearson’s r) between each stimulus pair was computed and aggregated in four representational dissimilarity matrices (RDMs); the obtained RDMs were normalized in a 0-1 range; finally, the four RDMs were averaged in the fixed model RDM, which was correlated to brain activity patterns in the subsequent analyses.

### Stimuli and behavioral segmentation of foreground and background

We selected from the 1870 images used by (Kay KN *et al.* 2008) a sub-sample of 334 pictorial stimuli which are also represented in the Berkeley Segmentation Dataset 500 (BSD) (Arbelaez P *et al.* 2011). For each BSD image, 5-7 subjects manually performed an individual “ground-truth” segmentation, which is provided by the authors of the dataset (http://www.eecs.berkeley.edu/Research/Projects/CS/vision/grouping/resources.html). Although figure-ground judgement is rather stable across subjects (Fowlkes CC *et al.* 2007), we selected the largest patch - manually labeled as foreground - among the 5-7 behavioral segmentations, in order to build a foreground binary mask (Figure S1). For each image, this mask was then down-sampled and applied to the original stimulus to isolate the foreground and the background pixels (Kay KN *et al.* 2008-http://crcns.org/data-sets/vc/vim-1).

### fMRI Data

The fMRI data used in this study are publicly available at http://crcns.org/data-sets/vc/vim1. Two subjects were acquired using the following MRI parameters: 4T INOVA MR, matrix size 64x64, TR 1s, TE 28ms, flip angle 20°, spatial resolution 2 × 2 × 2.5 mm^3^. For each subject five scanning sessions (7 runs each) were performed on five separate days. The stimuli were 1870 greyscale natural images with diameter 20° (500px), embedded in a grey background, and were presented for 1s, flickering at 5Hz, with an ISI of 3s. Subjects were asked to fixate a central white square of 0.2°(4px). Seven visual regions of interest (ROIs) - V1, V2, V3, V3A, V3B, V4 and LOC - were defined and brain activity related to stimulus presentation was extracted from these regions. For additional details on pre-processing, retinotopic mapping and ROIs localization, please refer to (Kay KN *et al.* 2008).

### Computational Models

In accordance with a previous fMRI study, we selected four well-assessed untrained computational models, which showed significant correlations with brain activity patterns in early visual areas as well as LOC (Khaligh-Razavi SM and N Kriegeskorte 2014). The four models comprise: GIST (Oliva A and A Torralba 2001), Dense SIFT (Lazebnik S et al. 2006), Pyramid Histograms of Gradients (PHOG) (Bosch A et al. 2007) and Local Binary Patterns (LBP) (Ojala T et al. 2001). For an exhaustive description of the four models – and links to Matlab codes – see the work by Khaligh-Razavi (2014) and Khaligh-Razavi and Kriegeskorte (2014).

### Representational Similarity Analysis (RSA)

For each filtered image, we collected feature vectors from the four computational models (GIST, PHOG, LBP and Dense SIFT), and RDMs were then obtained (1 minus the Pearson correlation metric). These four RDMs were normalized in a range between 0 and 1, and averaged to obtain the fixed biologically plausible model of the stimuli (for a graphical representation of the process, see Figure 2D). Single subject RDMs were similarly computed using fMRI activity patterns for each of the seven ROIs, and then averaged across the two subjects. We used Spearman’s rho (ρ) to assess the correlation between the RDM from each step of the image filtering procedures and the RDM of each brain ROI. For each correlation value, the standard error was estimated with bootstrapping of the stimuli –1,000 iterations (Efron B and R Tibshirani 1993).

In addition, as each ROI may show a distinct signal-to-noise ratio, we computed a noise estimation by correlating the brain RDMs extracted from the two subjects. This estimation of ROI-specific noise was used to normalize the correlation coefficients (normalized Spearman’s rho, Nρ). The same normalization procedure has been already employed in fMRI studies based on voxelwise encoding (Huth AG et al. 2016).

### Foreground enhancement testing

A permutation test was performed to statistically assess the enhancement of the information retained in the behavioral segmented foreground, and to rule out the possible confound due to a “fovea-to-periphery” bias that characterizes natural images (Figure S2). For each iteration of this procedure, the 334 foreground masks were shuffled and a random foreground segmentation was associated to each stimulus. Of note, this set of randomly-segmented images had the same distribution of masked portions of the visual field as the one from the behavioral segmentation, so the same amount of information was isolated at each permutation step. This procedure was repeated 1,000 times, to build a null distribution of alternative segmentations: four examples of random segmentation are shown in Figure 2B. For each permutation step, features were extracted from each randomly segmented image and RSA was performed using the procedure described above.

### Parametric filtering procedures

In order to investigate differential processing of foreground and background in the visual system, we employed three different filtering procedures (contrast - through alpha channel modulation - low- and high-pass filtering of spatial frequencies) applied parametrically (99 steps each) to the foreground or the background. For each filtering procedure, three examples of manipulated images are represented in Figure 2C. For low- and high-pass filtering, we employed a Butterworth filter (5^th^ order), linearly sampling from a log-transformed distribution of frequencies ranging from 0.05 to 25 cyc/°, while keeping the root mean squared (RMS) contrast fixed.

### Background suppression testing

To test background suppression, we performed three separate Welch t-tests. For contrast, low-pass and high-pass filtering we compared the maximum Nρ obtained by filtering out (i.e., suppressing) the background information with the Nρ value corresponding to the same filtering step obtained by suppressing the foreground information (p < 0.05, one-tailed). The three resulting p-values were aggregated using the Z-transform method. In this way, we tested whether applying the optimal filtering level for the background (ranging from intact to completely suppressed) to the foreground part of the image would lead to a lower correlation value with brain activity, thus revealing a specific mechanism of background rather than foreground suppression.

### Gaussian filtering of the foreground masks

In order to test to what extent the exactness of foreground borders explains the similarity between the isolated foreground mask and brain activity, an additional filtering procedure was computed. The selected behavioral masks were processed using a parametric Gaussian filter, whose sigma increased by 2 pixels at each step while keeping the segmented area constant. Therefore, the resulting masks provided an approximate segmentation of the foreground figure depending on the sigma of the filter. These masks were then applied to the original stimuli, and for each of these steps the correlation with fMRI activity patterns was computed. Three examples of this procedure are represented in Figure 2C and the results are shown in Figure 5G.

### Neural images

For each ROI, the effects of the filtering procedures were then combined, to build “neural images”. To this aim we used the filtering step with the highest correlation between the fixed model and brain activity, for foreground and background. In detail, we averaged the best images for the low- and high-pass filters, and multiplied each pixel for the preferred alpha-channel value (contrast). Lastly, the foreground mask employed for the neural images was chosen as the best step in Gaussian filtering procedure described above.

### Significance testing

To asses the statistical significance of the correlations obtained with RSA in all the above mentioned filtering procedures, we built a robust ROI-specific permutation test (1,000 iterations), by randomly sampling voxels of the occipital lobe not located in any of the seven ROIs. We labeled these voxels as ‘control-voxels’. This procedure has the advantage to be resilient to biases in fMRI data (Schreiber K and B Krekelberg 2013), instead of simply taking into account the distribution of the RDM values, as in Khaligh-Razavi S-M and N Kriegeskorte (2014). In addition, the procedure that we developed is also useful to control for the effects related to number of voxels and to the signal-to-noise of each ROI.

First, for each ROI we computed the standard error of the ROI-specific noise estimation with bootstrap resampling of the stimuli (1,000 iterations). Second, a number of control voxels equal to the number of voxels was randomly selected within each ROI, and the activity of these control voxels in response to the stimuli were used to build a null RDM. Third, the correlation between the null RDMs of the two subjects was computed. However, since we aimed at matching the signal-to-noise ratio of the null distribution to that of each ROI, the null RDM was counted as a valid permutation only if the single subject RDMs correlated to each other within a specific range (i.e., ROI-specific noise estimation ± standard error). Finally, for each step of the filtering procedures, each of the 1,000 ROI-specific null RDMs were correlated with the fixed model RDM to obtain a null distribution of 1,000 Nρ values. A signed-rank test was used to assess the significance of the Nρ of the fixed model with brain activity. For each ROI, we controlled for multiple comparisons (696 tests), using the False Discovery Rate procedure (Benjamini Y and Y Hochberg 1995).

All analyses have been performed using Matlab (The Mathworks Inc.).

## Results

### Comparison of intact and behaviorally segmented images

Three fixed descriptions of the stimuli were created (Figure 2A). RSA results show a significant correlation (p < 0.05 FDR corrected) between the intact description of images and brain activity in all the ROIs but V3A (V1: Nρ = 0.65; V2: Nρ = 0.74; V3: Nρ = 0.57; V3B: Nρ = 0.54; V4: Nρ = 0.51; LOC: Nρ = 0.59). Similarly, the segmented foreground RDM shows a significant correlation in all the ROIs, but V3 and V3A (V1: Nρ = 0.26; V2: Nρ = 0.34; V3B: Nρ = 0.52; V4: Nρ = 0.59; LOC: Nρ = 0.64), while the segmented background achieves significant correlations in V1 and V2 only (V1: Nρ = 0.31; V2: Nρ = 0.42).

### Foreground Enhancement

To test whether the behavioral foreground segmentation from BSD was more tied to brain activity as compared to alternate configurations obtained by shuffling the segmentation patterns across stimuli (Figure 2B), we performed a specific analysis based on a permutation test.

As depicted in Figure 3B, the correct foreground configuration yielded a significantly higher correlation as compared to the examples from the shuffled dataset, thus suggesting that the enhancement of foreground-related information occurs during passive perception of natural stimuli in all the tested ROIs (V1: p = 0.002; V2: p < 0.001; V3: p < 0.001; V3A: p < 0.001; V3B: p < 0.001; V4: p < 0.001; LOC: p < 0.001).

**Figure 3.**
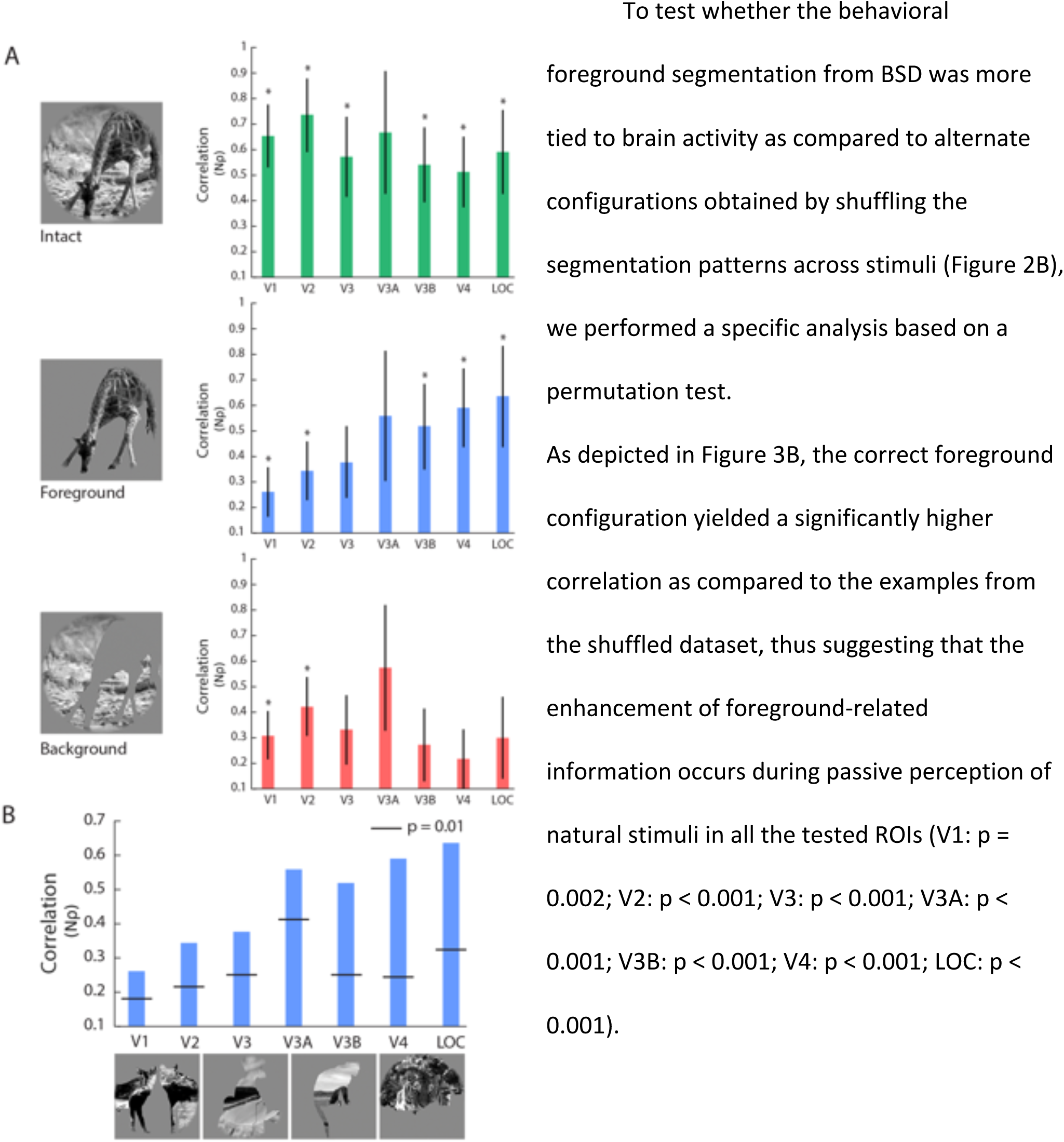
Foreground Enhancement in the Human Visual System. A) Correlation between the intact and segmented versions of the images and brain activity (* = p < 0.05, FDR corrected). Bars represent the standard error estimated with bootstrapping. B) To test foreground enhancement and rule out a “fovea-to-periphery” bias, the behavioral segmentation was tested against a null distribution of shuffled masks. All the tested ROIs yielded a significant correlation (p < 0.01, permutation test).

In addition, this analysis rules out a potential confounding effect related to a “fovea-to-periphery bias” in our image set - depicted in Figure S2. In fact, as already observed in literature, natural images are typically characterized by objects located at the center of the scene - see for instance the object location bias represented in Figure 3B in (Alexe B et al. 2010). However, since the spatial distribution and number of pixels were kept constant at each permutation step, we replicated the same “fovea-to-periphery bias” in the null distribution. Thus, we reject the possibility that foreground enhancement is driven by differences between the representation of fovea and periphery across the image set.

### Filtering procedures

As the correlation between the background RDM and brain activity is significant in V1 and V2 only (Figure 3A), we hypothesized that background-related information is suppressed in “higher” visual cortices. Notably, Poort and colleagues (2016) described background suppression as a different, but associated, phenomenon with respect to foreground enhancement. Thus, in order to better characterize where and how background suppression occurs in humans attending to natural images, a further analysis was performed by parametrically filtering out the background of each image, varying its contrast or spatial frequencies (low- and high-pass filtering; Figure 2C). RSA results for the parametric filtering approach are depicted in Figure 4. The correlation between manipulated images and the activity of V3A is not significant for any of the filtering steps, thus is not further discussed. Of note, this result is in agreement with the lack of significance for the correlation between the intact version of the image and brain activity of this region.

**Figure 4.**
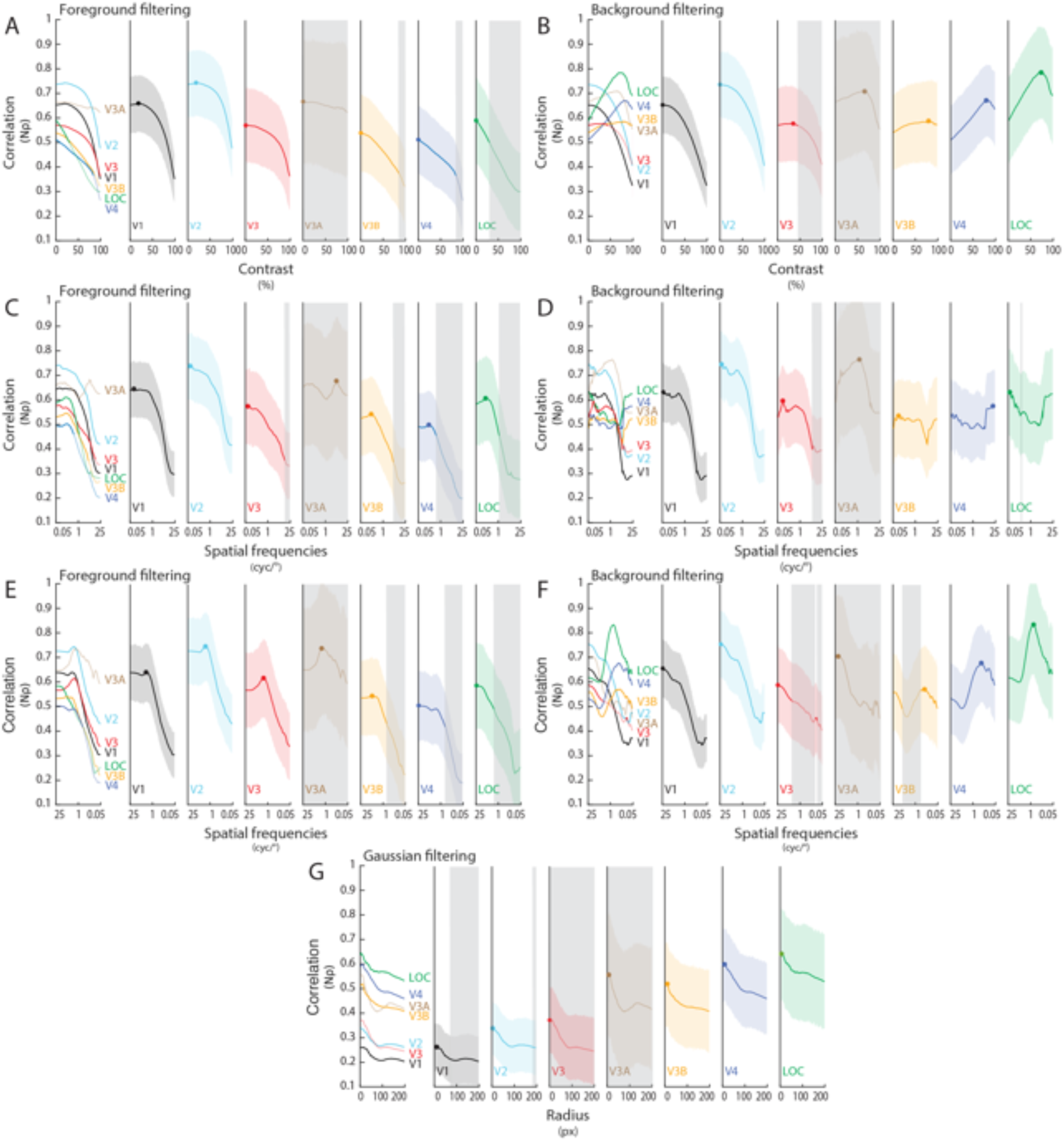
Results of the Filtering Procedures. Correlation between brain activity and contrast, high- and low-pass filtering applied to the foreground (A, C, E) and to the background (B, D, F). Panel G shows the correlation pattern for the Gaussian filter. Gray regions mark not significant correlations (p > 0.05, FDR corrected) while colored shaded areas represent the standard error estimates.

For the contrast filtering procedure applied to the background, the activity of V3 is significantly associated to degrees of filtering lower than 43%. All the other ROIs retain significant correlations for all the filtering steps. The maximum correlation with brain activity in each ROI is reached at the filtering degree of: 0% in V1; 0% in V2; 35% in V3; 80% in V3B; 80% in V4; 74% in LOC.

When the same procedure is applied to the foreground, the correlation is significant for degrees of filtering lower than 83% in V3B, 82% in V4 and 28% in LOC. All the other ROIs retain significant correlations for all the filtering steps. The maximum correlation with brain activity is reached at the filtering degree of: 19% in V1; 19% in V2; 0% in V3; 0% in V3B; 0% in V4; 0% in LOC.

For the high-pass filtering procedure applied to the background, correlation in V3 is significant until spatial frequencies higher than 6.6 cyc/° are filtered out. Correlation in LOC is always significant but at a single step, when frequencies between 0.26-0.28 cyc/° are filtered out. For all the other ROIs correlation is always significant. The maximum correlation with brain activity in each ROI is obtained when spatial frequencies in the range between 25 cyc/° and 0.05 cyc/° in V1, 0.06 cyc/° in V2, 0.10 cyc/° in V3, 0.11 cyc/° in V3B, 19.37 cyc/° in V4, 0.51 cyc/° in LOC are retained.

When the same procedure is applied to the foreground, the correlation is significant until spatial frequencies higher than 13.25 cyc/° in V3, 4.81 cyc/° in V3B, 0.56 cyc/° in V4 and 1.20 cyc/°in LOC are filtered out. For all the other ROIs correlation is significant at each step. The maximum correlation with brain activity is obtained when spatial frequencies in the range between 25 and 0.84 cyc/° in V1, 0.69 cyc/° in V2, 0.69 cyc/° in V3, 0.22 cyc/° in V3B, 0.22 cyc/° in V4, 0.19 cyc/° in LOC are retained.

For the low-pass filtering procedure applied to the background, correlation in V3 is significant until spatial frequencies lower than 3.7 cyc/° are filtered out. An exception is represented by the correlation between the V3 RDM and the fixed model RDM that retains a significant value when only frequencies between 0.11- 0.12 cyc/° are included. Correlation in V3B is significant unless only frequencies in the 0.5 - 7.0 cyc/° range are included. For all the other ROIs correlation is always significant. The maximum correlation with brain activity is obtained when spatial frequencies in the range between 0.05 and 24.94 cyc/° in V1, 23.41 cyc/° in V2, 24.94 cyc/° in V3, 0.30 cyc/° in V3B, 0.34 cyc/° in V4, 0.72 cyc/° in LOC are retained.

When the same procedure is applied to the foreground, the correlation is significant until the spatial frequencies lower than 0.72 cyc/° in V3B, 0.72 cyc/° in V4 and 2.39 cyc/°in LOC are filtered out. For all the other ROIs correlation is always significant. The maximum correlation with brain activity is obtained when spatial frequencies in the range between 0.05 and 2.72 cyc/° in V1, 2.11 cyc/° in V2, 1.98 cyc/° in V3, 1.86 cyc/° in V3A, 5.13 cyc/° in V3B, 23.41 cyc/° in V4, 24.94 cyc/° in LOC are retained.

### Background Suppression

To test the significance of background suppression we compared, for each ROI and each filtering procedure, the maximum Nρ achieved by filtering the background against the correlation value for the same degree of filtering applied to the foreground. Results show a significant effect only in V4 and LOC, indicating that background suppression occurs only in these regions (V4: p < 0.01; LOC: p = 0.04).

### Exactness of foreground borders

To test to what extent the exactness of foreground borders explains the similarity between the BSD masks and brain activity, a Gaussian filter was parametrically applied to the foreground masks and the resulting correlation was measured for each ROI (Figure 4 G). The exact or nearly exact version of the foreground segmentation showed the maximum correlation with brain activity in all the ROIs (V1: max step = 6 out of 100; 12px sigma; V2: max step = 1 out of 100; 0px sigma; V3: max step = 1 out of 100; 0px sigma; V3A: max step = 1 out of 100; 0px sigma; V3B: max step = 1 out of 100; 0px sigma; V4: max step = 4 out of 100; 8px sigma; LOC: max step = 3 out of 100; 6px sigma). Specifically, the correlation with brain activity is significant for sigma values lower than 2.9° (72px) in V1 and lower than 7.3° (184px) in V2. For all the other ROIs the correlation is always significant.

## Discussion

In the present study, we illustrated how the manipulation of low-level properties of natural images, and the following correlation with brain responses during passive viewing of the intact stimuli, could disclose the behavior of different brain regions along the visual pathway.

Employing this pre-filtering modeling approach, we were able to collect three different evidence indicating that scene segmentation is an automatic process that occurs during passive perception in naturalistic conditions, even when individuals are not required to perform any particular tasks, or to focus on any specific aspect of images.

First, we demonstrated that the correlation of fMRI patterns with foreground-related information is significant in V1, V2, V3B, V4 and LOC, while background-related information is significant in V1 and V2 only.

Second, our analyses specifically found that foreground enhancement is present in all the selected visual ROIs, and that this effect is driven neither by the foreground extent, nor by its location in the visual field. Thus, indirect evidence of figure-ground modulation of natural images could be retrieved in the activity of multiple areas of the visual processing stream. This is consistent with a recent study, which reported that border-ownership of natural images cannot be resolved by single cells, but requires a population of cells in monkey V2 and V3 (Hesse JK and DY Tsao 2016).

Finally, a proof of segmentation can be represented by the significant suppression of background-related information in V4 and LOC. On the contrary, earlier regions across the visual stream - from V1 to V3 – have a uniform representation of the whole image, as evident at first glance in the obtained neural images (Figure 5). Overall, these results further support the idea that foreground enhancement and background suppression are distinct, but associated, processes involved in scene segmentation of natural images.

**Figure 5.**
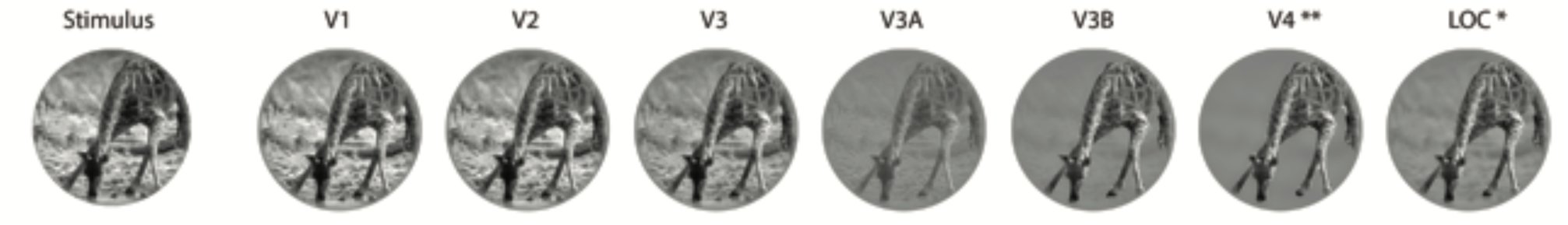
Neural Images Reveal Background Suppression in the Human Visual System. Neural images have been obtained as the combination of steps for different filtering procedures (contrast, Gaussian, low- and high-pass filtering), showing the highest correlation with brain activity for each ROI. Background suppression is significant in V4 and LOC (*: p < 0.05; **: p < 0.01).

### Foreground segmentation as a proxy for shape processing

The observed behavior of V4 and LOC is consistent with several investigations on shape features selectivity in these regions, and in their homologues in monkey (Carlson ET et al. 2011; Hung CC et al. 2012; Lescroart MD and I Biederman 2013; Vernon RJ et al. 2016). In fact, the extraction of shape properties requires segmentation (Lee TS *et al.* 1998), and presumably occurs in brain regions where background is already suppressed. Notably, “neural images” reconstructed from V4 and LOC are characterized by a strong background suppression, while the foreground is preserved. This is consistent with a previous neuropsychological observation: a bilateral lesion within area V4 led to longer response times in identifying overlapping figures (Leek EC et al. 2012). Hence, this region resulted to be crucial for accessing foreground-related computations, and presumably plays a role in matching the segmented image with stored semantic content in figure recognition. In accordance with this, a recent hypothesis suggests a role of V4 in higher-level visual functions, such as features integration or contour completion (Roe AW et al. 2012).

The preserved spatial resolution of foreground descriptive features (i.e., texture) in V4 and LOC – as shown in figures 4 and 5 - represents an additional noteworthy aspect that arises from our data. The progression from V1 towards higher-level regions of the cortical visual pathway is associated with a relative increase in receptive fields size (Dumoulin SO and BA Wandell 2008; Freeman J and EP Simoncelli 2011; Kay KN et al. 2015). However, it should be kept in mind that regions such as V4 demonstrate a complete representation of the contralateral visual hemifield, rather than selective responses to stimuli located above or below the horizontal meridian (Wandell BA and J Winawer 2011). The evidence that the foreground portion of “neural images” maintains fine-grained details in V4 and LOC seems to contrast the traditional view according to which these regions are more tuned to object shape (i.e., silhouettes), instead of being selective for the internal configuration of images (e.g. Malach R et al. 1995; Grill-Spector K et al. 1998; Moore C and SA Engel 2001; Stanley DA and N Rubin 2003). However, it has been shown that foveal and peri-foveal receptive fields of V4 do accomodate fine details of the visual field (Freeman J and EP Simoncelli 2011) and that the topographic representation of the central portion of this area is based on a direct sampling of the primary visual cortex retinotopic map (Motter BC 2009). Therefore, given the “fovea-to-periphery” bias found in our stimuli and in natural images, it is reasonable that an intact configuration of the foreground may be more tied to the activity of these brain regions, and that a richer representation of the salient part may overcome simplistic models of objects shape (i.e., silhouettes). Our result is also consistent with a recent study on monkeys that demonstrates a role of V4 in texture perception (Okazawa G et al. 2015).

Moreover, it is well known that selective attention represents one of the “active” cognitive mechanisms supporting figure segmentation (Qiu FT et al. 2007; Poort J *et al.* 2012), as suggested, for instance, by bistable perception phenomena (Sterzer P et al. 2009) or by various neuropsychological tests (e.g. De Renzi E et al. 1969; Bisiach E et al. 1976). In the present experiment, participants were asked to simply gaze a central fixation point without performing any overt or covert tasks related to the presented image. Nonetheless, we found evidence of a clear background suppression and foreground enhancement, suggesting that scene segmentation is mediated by an automatic process that may be driven either by bottom-up (e.g., low-level properties of the foreground configuration), or top-down (e.g., semantic knowledge) attentional mechanisms.

### Facing the challenge of explicit modeling in visual neuroscience

As predicting brain responses in ecological conditions is one the major goals of visual neuroscience, our study showed that the sensitivity of fMRI pattern analysis can represent an adequate tool to investigate complex phenomena through the richness of natural stimuli.

The standard approach in investigating visual processing in ecological conditions implies testing the correlation of brain responses from a wide range of natural stimuli with features extracted by different alternative computational models. This approach facilitates the comparison between the performances of competing models and could ultimately lead to the definition of a more plausible model of brain activity. However, the development of explicit computational models for many visual phenomena in ecological conditions is difficult, as testified by the extensive use of artificial stimuli in visual neuroscience (e.g. Carandini M et al. 2005; Wu MC *et al.* 2006).

Actually, even if computer vision is a major source of computational models and feature extractors, often its objectives hardly overlap with those of visual neuroscience. Computer scientists are mainly interested in solving single, distinct tasks (e.g., segmentation, recognition, etc.), while, from the neuroscientific side, the visual system is considered as a general-purpose system that could adapt itself to perform different behaviors (Medathati NVK et al. 2016). Consequently, while computer science typically employs solutions that rely only seldom on previous neuroscientific knowledge, and its goal is to maximize task accuracy (e.g., with deep learning), visual neuroscience somehow lacks of solid computational models and formal explanations, ending up with several arbitrary assumptions in modeling, especially for mid-level vision processing, such as scene segmentation or shape features extraction (for a definition see: Kubilius J et al. 2014).

In light of this, we believe that the manipulation of a wide set of natural images, and the computation of a fixed model based on low-level features, can offer a simple and biologically plausible tool to investigate brain activity related to higher-order computations. In fact, the results of this procedure can be depicted and are more intuitive as compared to descriptions obtained through formal modeling (Figure 5), thus highlighting interpretable differences rather than data predictions.

